# Self-organization of Long-lasting Human Endothelial Capillary Networks guided by DLP Bioprinting

**DOI:** 10.1101/2023.02.21.529380

**Authors:** Elsa Mazari-Arrighi, Matthieu Lépine, Dmitry Ayollo, Lionel Faivre, Jérôme Larghero, François Chatelain, Alexandra Fuchs

## Abstract

Tissue engineering holds great promise for regenerative medicine, drug discovery and as an alternative to animal models. However, as soon as the dimensions of engineered tissue exceed the diffusion limit of oxygen and nutriments, a necrotic core forms leading to irreversible damage. To overcome this constraint, the establishment of a functional perfusion network is essential and is a major challenge to be met. In this work, we explore a promising Digital Light Processing (DLP) bioprinting approach to encapsulate endothelial progenitor cells (EPCs) in 3D photopolymerized hydrogel scaffolds to guide them towards vascular network formation. We observed that EPCs encapsulated in the appropriate photopolymerized hydrogel can proliferate and self-organize within a few days into branched tubular structures with predefined geometry, forming capillary-like vascular tubes or trees of various diameters (in the range of 10 to 100 μm). Presenting a monolayer wall of endothelial cells strongly connected by tight junctions around a central lumen, these structures can be microinjected with fluorescent dye and are stable for several weeks *in vitro*. Interestingly, our technology has proven to be versatile in promoting the formation of vascular structures using a variety of vascular cell lines, including EPCs, human vascular endothelial cells (HUVECs) and human dermal lymphatic endothelial cells (HDLECs). We have also demonstrated that these vascular structures can be recovered and manipulated in an alginate patch without altering their shape or viability. This opens new opportunities for future applications, such as stacking these endothelial vascular structures with other cell sheets or multicellular constructs to yield bioengineered tissue with higher complexity and functionality.

## INTRODUCTION

Tissue engineering holds great promise in regenerative medicine, drug discovery and as an alternative to animal models. Although tremendous effort has been invested so far, engineered tissue development is confronted with a major challenge. Once the multicellular entity reaches a certain size, a necrotic core occurs due to the lack of an efficient perfusion system to sustain cell survival^1^. *In vivo*, blood vessels are essential for tissue homeostasis, supplying oxygen and nutrients, draining metabolic wastes and transporting immune cells. These functions are enabled by a highly ramified vascular network formed by an inner monolayer of endothelial cells (ECs), overlain either by smooth muscle cells (SMCs) in large and medium-sized vessels or by pericytes in capillaries.

For several years, many vascular tissue engineering strategies have been developed with the purpose of rebuilding or establishing a functional vascular system within an engineered tissue^2^. In this regard, the decellularization of donor arteries or veins is highly valued for its ability to generate acellular scaffolds from native tissue, which may optionally be recellularized prior to implantation^3,4^. However, the need to find a suitable donor limits its use. As a result, a completely different line of research based on the creation of vascularized organoids from basic components and cells has been widely pursued. Through the differentiation of human pluripotent stem cells (hPSCs) into EC and pericytes, Wimmer et al. developed a self-organizing 3D vascular organoid capable of forming a stable and perfused vascular tree when transplanted into immunocompromised mice^5^. Additionally, the differentiation of adult or induced stem cells has allowed the manufacture of various kinds of kidney, skin or brain organoids composed of a vascular-like network^6–8^. As demonstrated with the recent development of an organoid model of diabetic vasculopathy^9^, 3D culture and stem cell differentiation approaches offer a valuable alternative to animal models. Nevertheless, without appropriate spatial cues to guide cell self-organization this process remains highly stochastic, with no control over the size, accessibility and 3D geometry of the vessels formed in the bulk of a hydrogel.

To address this challenge, several engineering strategies offer various levels of control over these criteria by guiding spontaneous cell self-organization^10^, including sheet-based engineering^11–13^ and biocompatible polymer-scaffolds approaches^14–18^. Indeed, three-layered vascular vessels of various diameters were manufactured by rolling a single cell sheet comprising a polydimethylsiloxane (PDMS) membrane that served as a support for human ECs, SMCs and fibroblasts^19^. Recently, bilayer vascular vessels of various diameters were obtained by co-extrusion of a sacrificial polymer combined with hydrogels comprising human EC and SMCs^20^, and by seeding both cell types onto a synthetic bilayer heparinized tubular structure created by combining electrospinning and freeze-drying methods^21^. However, with a resolution ranging from ~100 μm to 2 mm in diameter, these technologies are unable to reproduce small vessels and more specifically capillaries (whose diameter varies from 20 μm down to a few micrometers). *In vivo*, capillaries form a complex and nevertheless well-organized meshed network that ensure the proper supply of oxygen and nutriments to all tissues. To this end, adjacent capillaries are separated from each other by a maximum distance of 200 μm^22^. In order to avoid the formation of an oxygen gradient, which might induce unforeseen cell differentiation or necrosis^1,23^, reproducing the capillary bed connected to a higher vascular tree therefore appears to be a key challenge for vascular tissue engineering.

Interestingly, digital light processing (DLP) is a promising 3D bioprinting technology that combines complete control over the 3D geometry at high resolution, while allowing high accessibility to the polymerized structure^24^. Using this approach, cells can be encapsulated in 3D photopolymerized hydrogels of the desired shape. By mimicking the mechanical properties of the natural cell environment^25^, the 3D photopolymerized hydrogel also provides spatial cues to guide both cell proliferation and polarization. Using this method, we have previously manufactured a branched tubular epithelial network successfully mimicking the intrahepatic biliary tree^26^.

In this study, we used human endothelial progenitor cells (EPCs) from cord blood and DLP bioprinting to produce vascular structures of controlled size and geometry by photopolymerization. After one week of culture, EPCs would self-organize into ramified vessels with a central lumen guided by the predefined geometry of the polymerized hydrogel. These branched vascular structures could be maintained for weeks in culture. Consisting of a monolayer of ECs connected by tight junctions, these vessels could be microinjected with a fluorescent dye and could be recovered and manipulated in a flexible alginate patch.

## MATERIAL AND METHODS

### Cell sources and reagents

Endothelial progenitor cells (EPCs) - EPCs were collected from two independent donors by isolating mononuclear cells from human umbilical cord blood according to the previously published protocol^27^. Briefly, mononuclear cells were isolated by density gradient centrifugation over Ficoll for 20 min at 500×g, washed three times in PBS and cultured on collagen-coated 6-well plates in endothelial growth medium (EGM-2, Lonza) composed of endothelial basal medium supplemented with 2% fetal bovine serum, 0.2 μg/mL hydrocortisone, 1 μg/mL ascorbic acid, 5 ng/mL human epidermal growth factor, 20 ng/mL of insulin-like growth factor, 22.5 μg/mL heparin, 0.5 ng/mL vascular endothelial growth factor, 10 ng/mL human fibroblast growth factor, and 2% gentamicin sulfate-amphotericin. After a few days in culture, non-adherent cells were removed and discarded and adherent EPCs were banked.

Human umbilical vein endothelial cells large volume (HUVECs XL) and human dermal microvascular endothelial cells (HDMECs) were purchased from Lonza and cultured in EGM-2 (Lonza) under 5% CO_2_ at 37 °C. Human dermal lymphatic endothelial cells (HDLECs) were purchased from Promocell and cultured in endothelial cell growth medium MV 2 (Promocell) under 5% CO_2_ at 37 °C.

For all cell sources, only cells from passages 2-6 were used in the following experiments. All reagents were obtained from Sigma Aldrich, France unless otherwise specified.

### Fabrication of hydrogel structures encapsulating ECs using DLP bioprinting

Photopolymerizable hydrogel formulations were prepared in PBS with final concentrations of 1% or 2% (w/v) methacrylated gelatin (G^MA^), 0.1%or 0.2% (w/v) type I methacrylated collagen (C^MA^, Advanced BioMatrix), 0.3% (w/v) methacrylated hyaluronic acid (HA^MA^, Advanced BioMatrix), 0.4% (w/v) fibrinogen (FG), and 0.5% (w/v) lithium phenyl-2,4,6 trimethyl-benzoyl phosphinate (LAP, Allevi). ECs were harvested with 0.25% trypsin-EDTA and the cell pellet was resuspended at a cell density of 6.10^6^ cells/mL directly in the photopolymerizable hydrogel formulation.

The DLP setup consisted of an Olympus IX83-inverted microscope (Olympus Corporation) equipped with a DMD-based UV patterned illumination device (PRIMO, Alveole), with a laser wavelength of 375 nm. Rasterized image files were sent to the DMD device, an array of approximately two million micromirrors that can be controlled individually to generate the optical pattern. The optical photopattern was projected through the 10x objective of the microscope. 35 μL of the photopolymerizable hydrogel formulation containing 6.10^6^ cells/mL were dispensed into the space between a methacrylated glass coverslip and a polydimethylsiloxane (PDMS) pad, separated by a silicone spacer of 250μm height (SIN 400 T, Sterne, France), and exposed to the UV pattern at a dose of 20 mJ/mm^2^ unless stated otherwise. After UV exposure, the PDMS stamp was peeled off and the structures were washed with PBS to remove the unpolymerized fraction of the hydrogel/cell formulation. The glass coverslip bearing the hydrogel/cell structures was transferred to a petri-dish or a well plate and cultured in EGM-2 culture medium. Culture medium was renewed 3 times a week.

### Cell viability assays

Each previously described photopolymerizable hydrogel mix was gently mixed with EPCs at a cell concentration of 6.10^6^ EPCs/mL and was photopolymerized with various UV doses: 10, 20, 40, and 60 mJ/mm^2^ doses. Cell viability was checked, 4 h and 24 h after photopolymerization respectively, by staining with a live and dead assay kit (Live/Dead Cell Viability Assay, Thermo Fisher Scientific) following the manufacturer’s manual.

### Fixation, immunofluorescent staining, and confocal microscopy

To investigate the vascular endothelium formation in vitro, encapsulated EPCs in photopolymerized hydrogel structures were cultured for a week in EGM2 culture medium and then fixed in 4% PFA in PBS for 15 min at room temperature, washed three times for 10 min in PBS and incubated for 1 h in blocking solution (5% normal goat serum, Cell Signaling Technology #5425, 0,3% Triton X-100 in PBS) at room temperature. The blocking was followed by overnight incubation with the primary antibodies in PBS with 1% BSA, 0,3% Triton X-100 at +4°. After three 10 min washings in PBS at room temperature, samples were incubated with secondary antibodies in PBS with 1% BSA, 0,3% Triton X-100 for 2 h at room temperature. Finally, samples were mounted using Aqua-Poly/Mount mounting medium and imaged using a confocal laser scanning microscope Zeiss LSM780 equipped with a 40× objective. In the following, is the list of used antibodies and dyes with the dilution ratios relative to stock concentrations recommended by the manufacturers: anti-CD31 (Abcam #ab28364) – 1:100; anti-ZO-1 (Invitrogen #33–9100) – 1:100; goat anti-Mouse IgG (Cy3) (Abcam #ab97035) – 1:200; goat anti-Rabbit IgG (Alexa Fluor 488) (Thermo Fisher Scientific #A-11034) – 1:200; phalloidin (Alexa Fluor 647) (Thermo Fisher Scientific #A22287) – 1:200; DAPI (Roche #10 236 276 001) – 4 μg/ml.

### Time-lapse image acquisitions

Time-lapse microscopy was performed at 37°C and 5% of CO_2_, with images taken at 30-min intervals using a Leica DMi8 microscope equipped with a 10x objective.

### Rheological analysis

The rheological features were evaluated for all three different photopolymerized hydrogel formulations: G^MA^/HA^MA^/FG *2%:0.3%:0.4% (w/v)*, C^MA^/HA^MA^/FG *0.2%:0.3%:0.4% (w/v)* and G^MA^/C^MA^/HA^MA^/FG *1%:0.1%:0.3%:0.4% (w/v)*

#### Oscillatory measurements

Storage and loss moduli of photopolymerized hydrogels were measured by a Discovery HR 2 rheometer (TA Instruments) equipped with a parallel plate geometry (d= 25 mm). Samples with a thickness of 250 μm and 25 mm in diameter were photopolymerized with either a 20 mJ/mm^2^ dose or a 40 mJ/mm^2^ dose. The photopolymerized samples were subjected to oscillatory measurements after being incubated in EGM-2 medium for 1 day at 37°C. The frequency sweeps were conducted at room temperature in the range of 0.1-10 Hz, and with a constant strain rate of 2%, considered to be in the linear viscoelastic range.

#### Compression tests

The elastic modulus of the resulting photopolymerized hydrogels was determined on a Microtester G2 (CellScale) on rectangular pads with a thickness of 250 μm and 6 mm^2^ area at room temperature. Before testing, the samples were incubated for 1 day in EGM-2 at 37 °C to reach swelling equilibrium. The compression test was conducted at 10-20% strain with 2 μm/s strain rate. The elastic modulus of each sample was calculated from the linear region of the stress-strain curve using Origin software with the force and displacement data collected from the Cell Scale software.

### Microinjection setup

The micropipette used for the microinjection of vascular tubes was made from borosilicate glass tubes with an outer diameter of 1.2 mm and an inner diameter of 0.94 mm (Harvard Apparatus). The injection micropipette was pulled using a micropipette puller (PC-100, NARISHIGE). The resulting injection micropipette was backfilled with 5-10 μL of 500 μg/mL fluorescein solution and was mounted on the microinjection manipulator (InjectMan 4, Eppendorf) connected to a pneumatic microinjection pump (FemtoJet 4i, Eppendorf) with a compensation pressure of 35 hPa. The injection micropipette was introduced in the lumen on one end of the targeted tubular network while the distal end was cut using surgical scissors. Once the tip of the micropipette was positioned within the lumen of the VSPPs, the fluorescein solution was injected at 150 hPa during 1 second and was tracked under an inverted microscope (Nikon Ti2, Nikon) using a 10X objective.

### Encapsulation of vascular networks within an alginate layer

After removing the culture medium, a drop of 50 μL of 1.5% Na-alginate (Wako Pure Chemical Industries) was placed over the vascular structures on the glass coverslip inside a 1 cm2 frame cut out from a 250 μm thick silicone sheet (SIN 400 T, Sterne, France). This allowed the drop to spread out evenly in a thin film over the structures. The glass coverslip was then gently immersed into a mixture of 100 mM CaCl_2_ and 3% w/w sucrose solution for 10 minutes. The CaCl_2_/sucrose solution was removed and replaced by EGM-2 culture medium. The alginate layer spontaneously detached from the glass coverslip after 15-30 minutes, and was transferred into a 35 mm petri dish and cultured in the EGM-2 under 5% CO_2_ at 37 °C.

## RESULTS AND DISCUSSION

ECs are known to spontaneously self-organize into “cord-like” structures when embedded in various hydrogel environments including collagen, gelatin, fibrin, and matrigel^28–31^. This self-organization is highly disorganized, branching happens randomly, and structures rarely form stable lumens or vascular networks which could subsequently be manipulated and integrated into a bioengineered tissue construct. We thus developed a technique using DLP bioprinting^24^ to control the microenvironmental cues necessary to promote the self-organization of endothelial progenitor cells (EPCs) into branched tubes with various size and shape (Fig 1A) using a similar experimental process described in^26^. Briefly, a photopolymerizable mix of hydrogels, cells and photoinitiator was placed between a glass coverslip and a PDMS roof, and UV light was projected as a DMD-generated, mozaic image through the 10x objective of a microscope. Non-polymerized hydrogel and cells were washed away and the resulting 3D structures were placed in the cell medium and cultured in a standard cell incubator for the indicated timeframes. Various DMD-generated images and the resulting structures at day 0 and day 2 are shown in Fig 1B. These structures include a line, with the aim of creating a single tube with a given diameter; a tree, to explore efficiency of creating a branching network as seen during angiogenesis; and a stylized capillary bed with line diameters going from 10 μm to 200 μm. Zoomed out images of the resulting constructs showing structures covering areas of 5×5mm approximately are shown Fig 1C.

**Figure 1:**
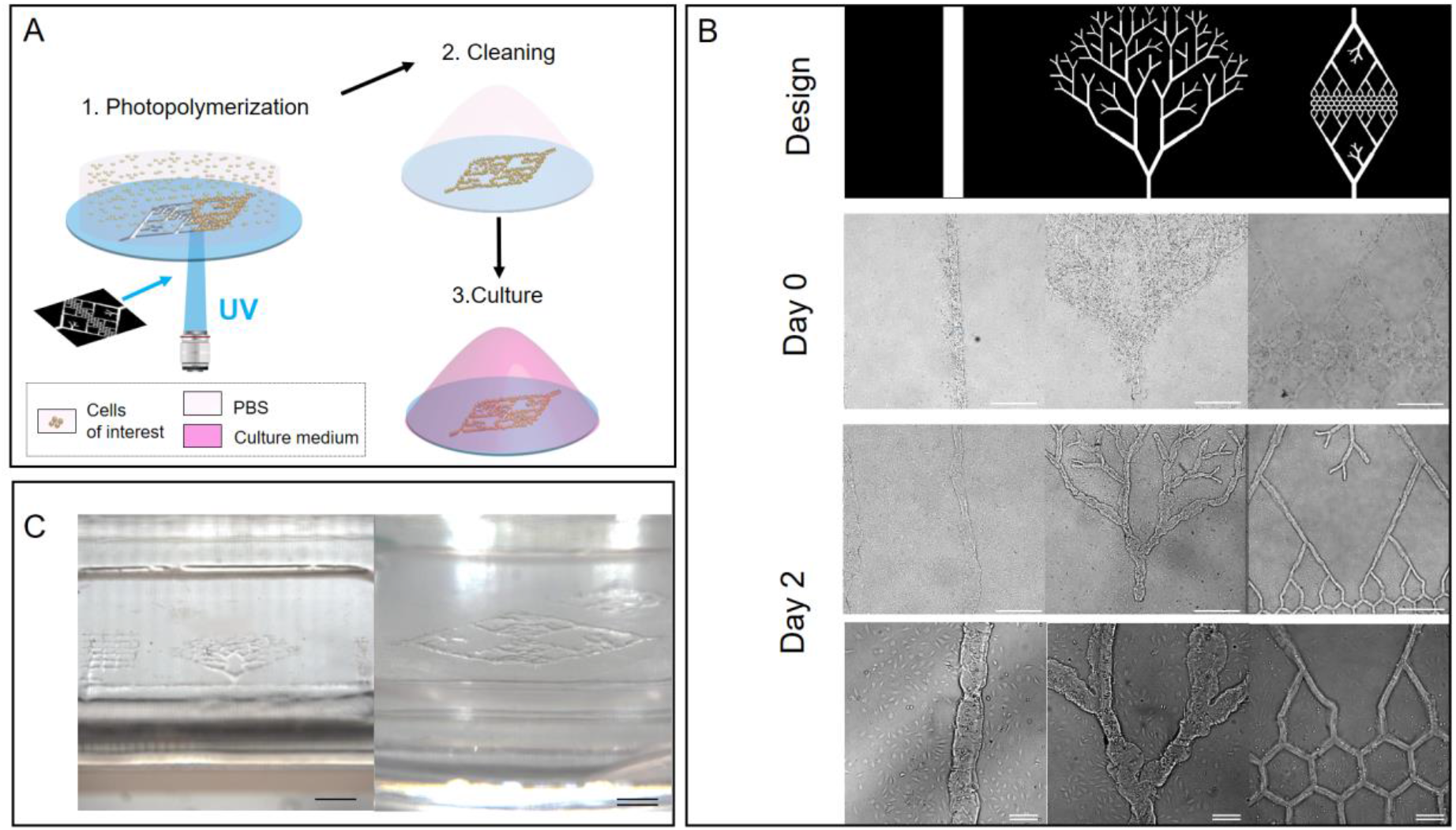
DLP photopolymerization of G^MA^C^MA^HA^MA^FG hydrogel structures encapsulating EPCs (A) Principle of DLP bioprinting. (B) Various digital patterns used to create photopolymerized structures embedding EPCs: a line, a branching tree and a capillary bed-type design, just after fabrication and 2 days after fabrication (C) Sagittal view of EPC photopolymerized structures after 7 days of culture. Scale bars are 500 μm and double scale bars are 50 μm. (C) Sagittal view of EPC photopolymerized structures after 7 days of culture. Scale bar is 2 mm and double scale bar is 1 mm.

### A four-component photopolymerizable hydrogel promotes the proliferation, migration, and self-organization of endothelial progenitor cells (EPCs) into stable branched tubuloid networks with predefined size and geometry

Based on our previous success with epithelial biliary cells^26^ and on published studies using endothelial cells^32^, we first explored various formulations incorporating either methacrylated gelatin (G^MA^) or methacrylated type I collagen (C^MA^) supplemented with fibrinogen (FG), methacrylated hyaluronic acid (HA^MA^), and lithium phenyl-2,4,6 trimethyl-benzoyl phosphinate (LAP) – a cytocompatible photoinitiator^33^ (Fig 1B Fig S1). Indeed, G^MA^ and C^MA^ are photopolymerizable hydrogels produced from gelatin and collagen respectively which sustain good cell viability following encapsulation and retain natural cell-binding motifs^34–36^. HA^MA^ is also a photopolymerizable hydrogel^37,38^ derived from hyaluronic acid, which is found ubiquitously in native tissues and plays an important role in many cellular responses, such as cell signaling or cell proliferation. FG is a component of the ECM that mediates cellular functions such as adhesion, spreading, proliferation, and migration of a variety of cell types, including fibroblasts, endothelial and epithelial cells. We observed that Endothelial Progenitor Cells (EPCs) encapsulated in either G^MA^/HA^MA^/FG (2%:0.3%:0.4% in w/v) or C^MA^/HA^MA^/FG (0.2%:0.3%:0.4% in w/v) formulations showed active proliferation, migration and reorganization into tubuloid structures over the first few days in culture (see movies M1, M2 and M3 in supplemental data). However, these observations lacked reproducibility and stability resulting in some cases in the rapid degradation and loss of the tubuloid structures, while the rate of success also appeared to be EPC donor-dependent (as observed in supplemental Fig S1).

The same hydrogel structures showed excellent stability over time in the absence of cells, indicating that the lack of stability was directly linked to the presence of living cells. We thus hypothesized that the gel structures were not strong enough to resist degradation by endothelial cells in contrast to what we had observed in a previous study with epithelial cells^26^, and tested other means to increase the stability of our formulations.

In a first approach, we increased the UV dose to increase chemical crosslinking and this indeed did lead to higher storage and loss moduli of the polymerized structures (Fig S2).

However, we also observed increased cell death in these samples (Fig S3) at day 1, from 60-70% viability at 10-20 mJ/mm^2^ down to 50-55% at 40 mJ/mm^2^, which is probably due to increased oxidative stress from the higher UV doses.

In a second approach, we screened higher concentrations of each hydrogel component, but were confronted to higher viscosity, heterogeneity, and less efficient washing of the structures after photopolymerization (data not shown). Finally, we studied a four-component mix based on existing successful studies using a hybrid mix of both gelatin and collagen with endothelial cells^39^. We thus mixed one volume of the gelatin formulation *G*^MA^/*HA*^MA^*/FG* with one volume of the collagen formulation *C*^MA^/*HA*^MA^/*FG* thus yielding the 4-component formulation: G^MA^/C^MA^/HA^MA^/FG 1%:0.1%:0.3%:0.4% in w/v. We expected that given both 3-component formulations exhibited comparable G’, G” and Y modulus (Fig 2A and B), the 4-component mix would present similar mechanical properties. Surprisingly, this was not confirmed experimentally, and the four-component mix showed a significant increase in both storage and loss modulus (Fig 2A), to reach values reported by Davidov et al.^40^ where they prepared a ECM matrix from decellularized arterial tissue which showed higher efficiency in generating stable angiogenic-type growth in the standard in vitro assay. This significant increase in terms of stiffness was confirmed in compression tests with nearly a two-fold increase in Young modulus from approximately 2 kPa for the 3-component hydrogels to 4 kPa for the 4-component hydrogel (Fig 2B), also consistent with values reported by others^41,42^.

**Figure 2:**
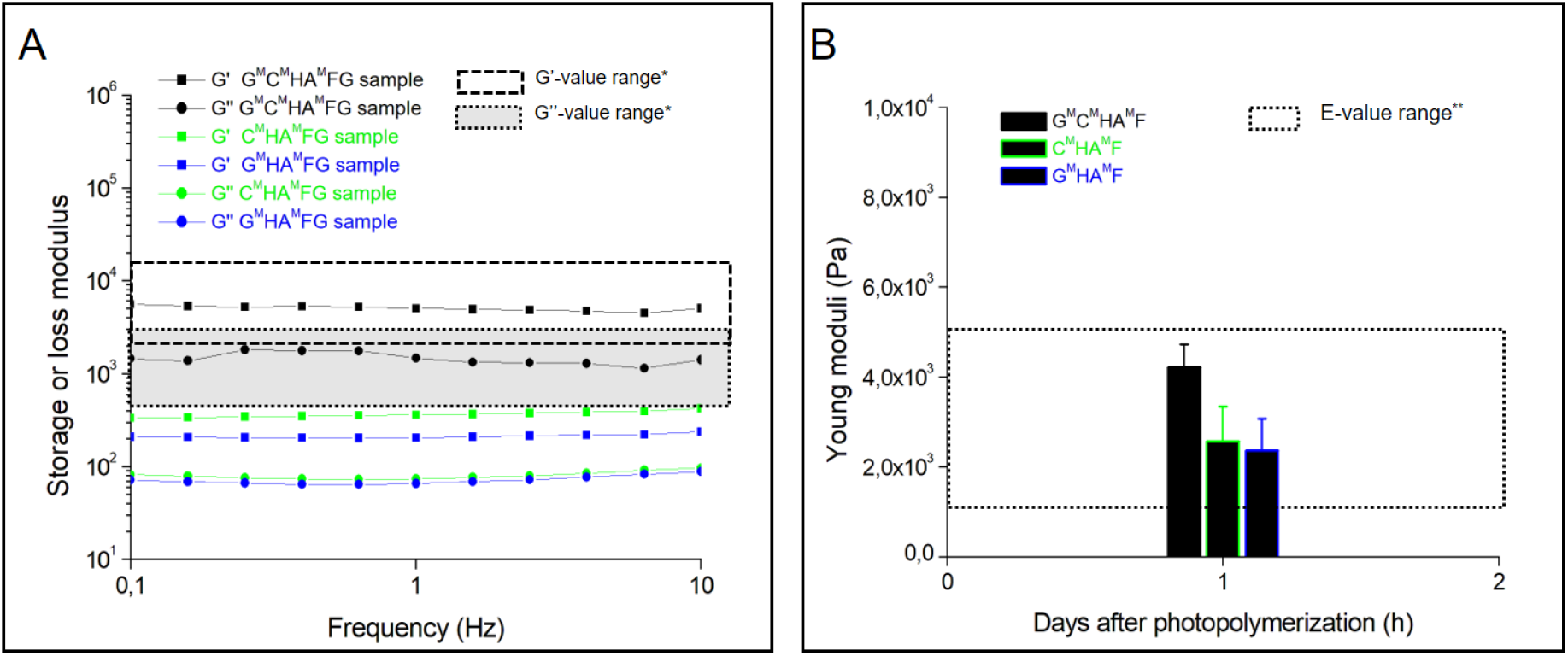
Rheological characterizations of photopolymerized 3- and 4-component hydrogel structures using either frequency-dependent oscillatory rheological analysis (A) or compression tests (B). *,**:Boxed areas correspond to modulus values of related ECM matrices published in (A) reference^40^ and in (B) references^41,42^.

We also observed that the four-component G^MA^/C^MA^/HA^MA^/FG hydrogel supported higher cell viability during and after the photopolymerization process compared to either G^MA^/HA^MA^/FG or C^MA^/HA^MA^/FG (Fig S3). More importantly, this hydrogel formulation showed substantially higher efficiency, reproducibility and stability over time in forming tubuloids with considerable migration and reorganization of the EPCs within them (Fig 1 and 3), and independently of the EPC donor (Fig S1).

**Figure 3:**
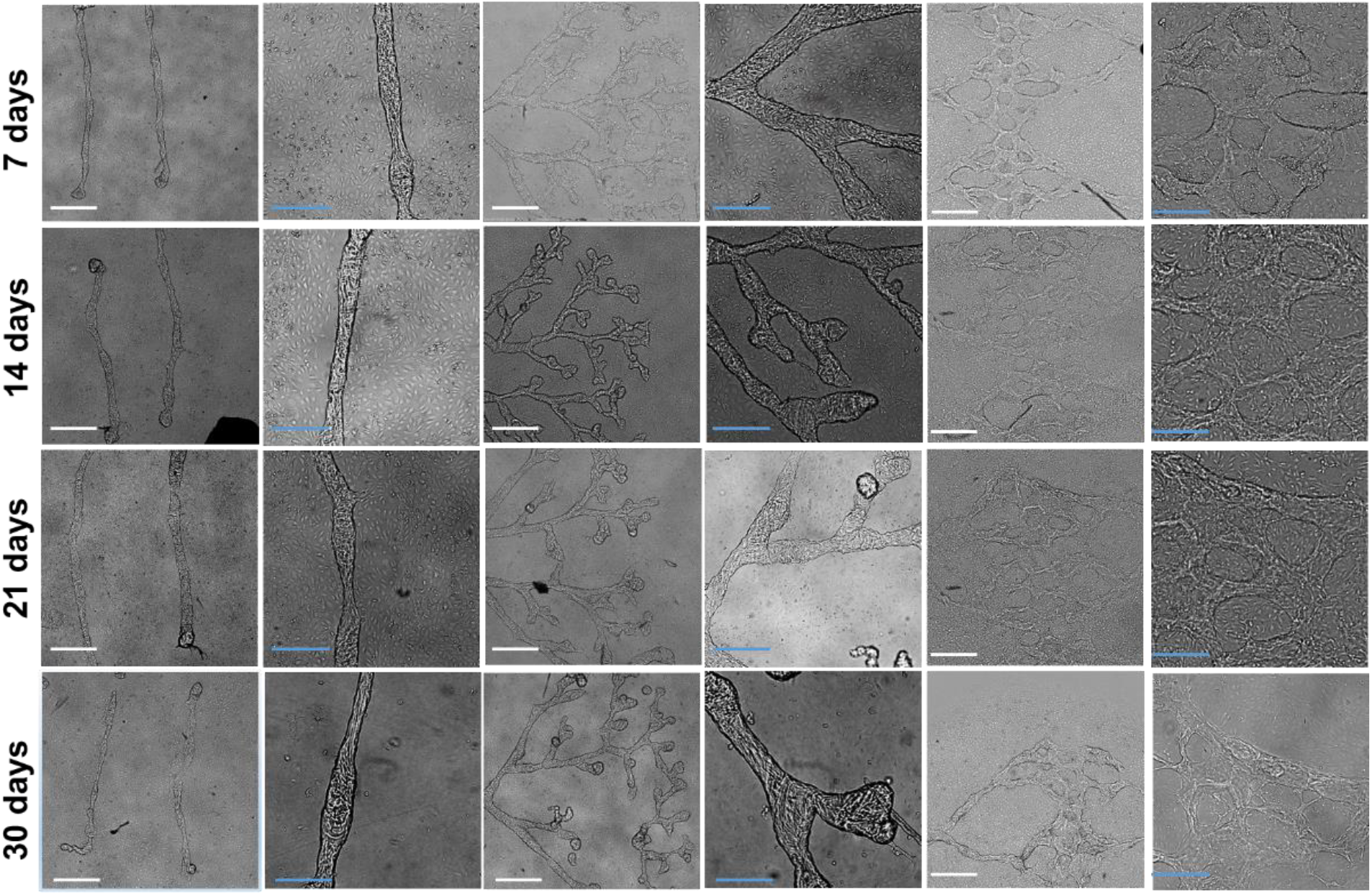
EPC photopolymerized hydrogel structures over time (between day 7 and day 30) Phase contrast images showing structures photo-polymerized through a 10X objective at 20 mJ/mm^2^. Hydrogel formulations are composed of G^MA^/C^MA^/HA^MA^/FG 1%:0.1%:0.3%:0.4% in w/v with 0.5% (wt/vol) photoinitiator. Blue scale bars are 200 μm and white scale bars are 500 μm.

Finally, this mix was tested successfully with other types of ECs, such as commercially-available human umbilical, dermal as well as lymphatic endothelial cells (HUVECs, HDMECs, and primary HDLECs, respectively). As shown in figures 3 and S4, all these commonly used sources of endothelial cells self-organized with similar overall behavior and kinetics to form tubuloid-type bodies of a wide diversity of shapes, branching and sizes. Line structures from 10μm to 200μm in width and millimeters in length thus generated tubuloid structures of approximately the same width and length, whereas branched trees and honeycombed patterns (Fig 1 and Fig 3) generated stable interwoven networks with good fidelity to the original design, spanning tens of mm^2^ (Fig 1B-C and Fig 3). The tree pattern was designed to mimic a developing capillary network, similar to what is observed during angiogenesis, whereas the honeycombed pattern with coalescing branches towards a single inlet and outlet on each side recapitulates the geometry of a stable capillary bed perfusing a tissue. All these tubuloid networks appeared to fully form and stabilize in approximately 5-7 days and then remained stable in *in vitro* culture for up to 21 days and more. (Fig 3). In the following sections, we pursued the study using primary EPCs from single donors as representative of the most challenging source of progenitor primary endothelial cells.

### After 7 days in culture, tubuloid structures developed a tight monolayer wall of endothelial cells spanning a central single lumen, mimicking the cell organization of a blood capillary

Confocal microscopy was used to characterize the internal cell organization of the tubuloid structures (Fig. 4). Labelling of F-actin, nuclei, tight junctions, and of CD31, specific for endothelial cells, showed that these tubuloid structures were exquisitely mimicking capillary endothelial tubes with a tight monolayer of cells forming a circumferential wall around a central lumen. In certain areas, this tube was observed anchored at a point on the glass coverslip (depicted in Fig4A with white arrows) and then rising a few tens of microns above the glass coverslip plane on which a standard 2D layer of endothelial cells was also visible. Intense ZO1 and CD31 labeling largely concentrated at the junctions between adjacent cells and showed no defects in the wall. As depicted in Fig. 4C, branching points imaged from tree-designs also exhibited a full external monolayer wall with a continuous Y shaped lumen. The lumen were approximately the same diameters as the original photopolymerized hydrogel structures as seen in Fig 4A and Fig 4B which show respectively tubes obtained with 10 μm and 50 μm hydrogel lines. The faint fluorescent signal observed inside the lumen was attributed to the typical autofluorescence of collagen (indicated by *), demonstrating that the initial hydrogel remained present to a certain extent. Interesting, darker patches inside the lumen could also be observed, sometimes spanning hundreds of microns over half the section of the tube (Fig 4B). This could therefore point to a partial degradation of this hydrogel scaffold over time. Interestingly, in these areas, the single monolayer cell wall did not appear to collapse, suggesting a different type of support, either linked to some osmotic pressure or the presence of newly synthesized extracellular matrix which would need further in-depth studies to elucidate.

**Figure 4:**
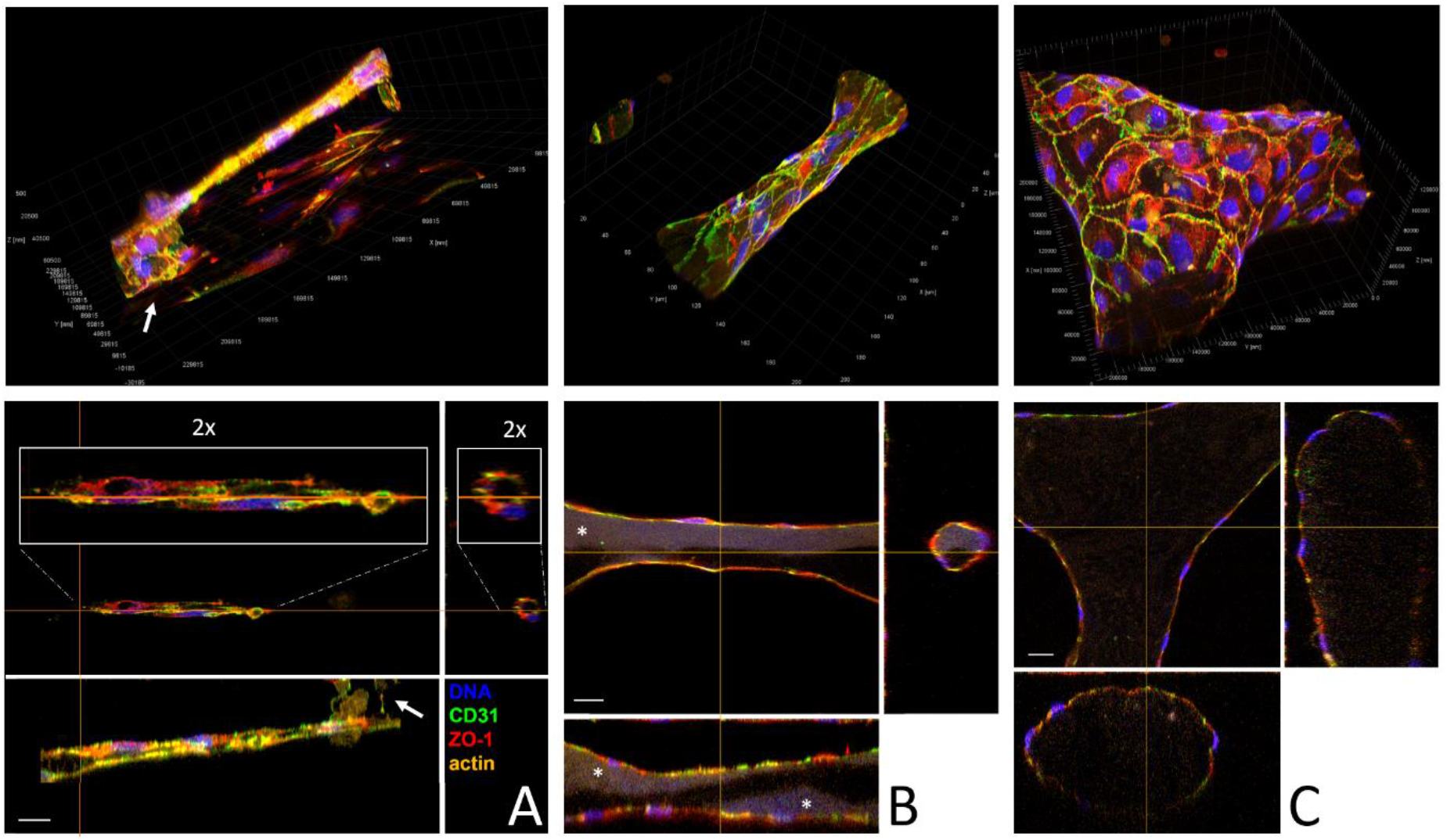
Characterization of EPC photopolymerized structures using confocal microscopy (day 7). Photo-polymerized hydrogels are composed of the same 4-component mix as above. Confocal images and orthogonal views show structures photo-polymerized through a 10X objective at 20 mJ/mm^2^ after 7 days of culture in (A) a 10μm line, (B) a 50μm line and in (C) at a branching point of a tree. Autofluorescence of the collagen is indicated with a * and anchoring points of the 10μm tube onto the glass slide is indicated with white arrows. 2x inserts correspond to zoomed areas on the sections of the 10 μm tube. Scale bar is 20 μm.

### Endothelial tubes can be microinjected with a fluorescent dye

A pulled glass capillary tip linked to a standard microinjection setup was positioned in the lumen of a seven-day-old branched vascular network or tube as depicted in Fig 5 under a baseline positive pressure of 35 hPa to avoid clogging the pipette. An injection at 150 hPa during one second was then applied to inject the fluorescein inside the luminal space. We found that fluorescence could be detected in the vascular tube only if the tube was physically opened at a distal end (outside the camera view). In this case, despite a high fluorescence background linked to the fluorescein dye leaking out, a high level of fluorescence is observed inside the lumen and fluorescein rapidly diffuses along the tubular network in less than a minute as illustrated in Fig 5, demonstrating that the lumen is uninterrupted even across multiple branching points.

**Figure 5:**
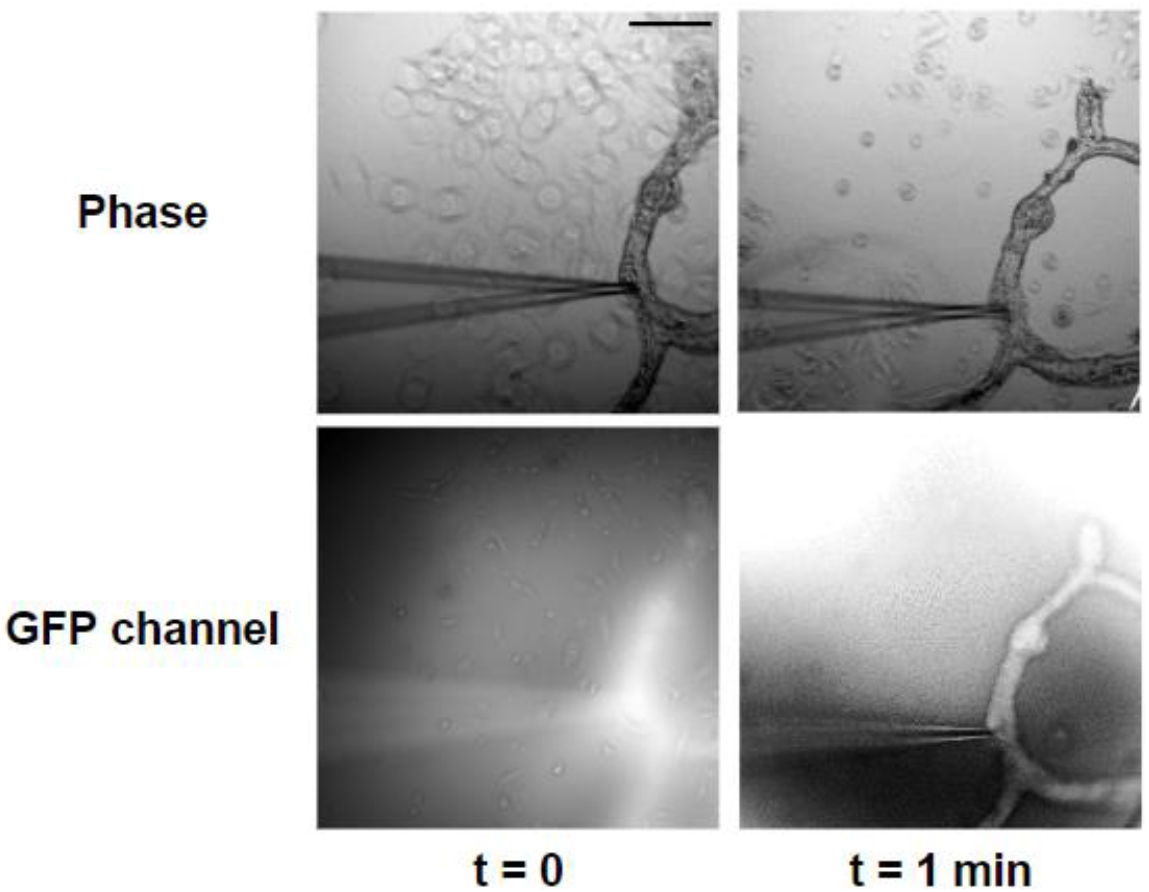
Microinjection of EPC tubes at 7 days of culture. Before perfusion, the microinjection pipet was inserted into the endothelial tube while the opposite side of the EPC tube was cut. Scale bar is 100 μm.

### Endothelial networks can be removed from the coverslip surface and manipulated as a flexible soft cell sheet for further tissue engineering applications

With the aim of integrating these highly organized endothelial networks with other bioengineered cell constructs and in particular with the aim of creating transplantable tissue, it became necessary to develop a method to remove the structures from the supporting glass coverslip without damaging them. As embedding cells in alginate hydrogels (as beads, fibers or sheets) is a common strategy to preserve cell function and viability^43–45^ while offering a convenient matrix for further manual handling, we explored various approaches and found that depositing a drop of 1.5%alginate inside a silicone frame surrounding the construct and then letting it gel by immersing it into a calcium solution was efficient in embedding the vascular structures in a thin layer. After transfer into cell culture medium, the thin sheet of alginate spontaneously delaminated from the glass chip, peeling away the vascular structures and thus providing an easy and reproducible way to recover and manipulate mature vascular structures while preserving their shape and viability. (Fig 6).

**Figure 6:**
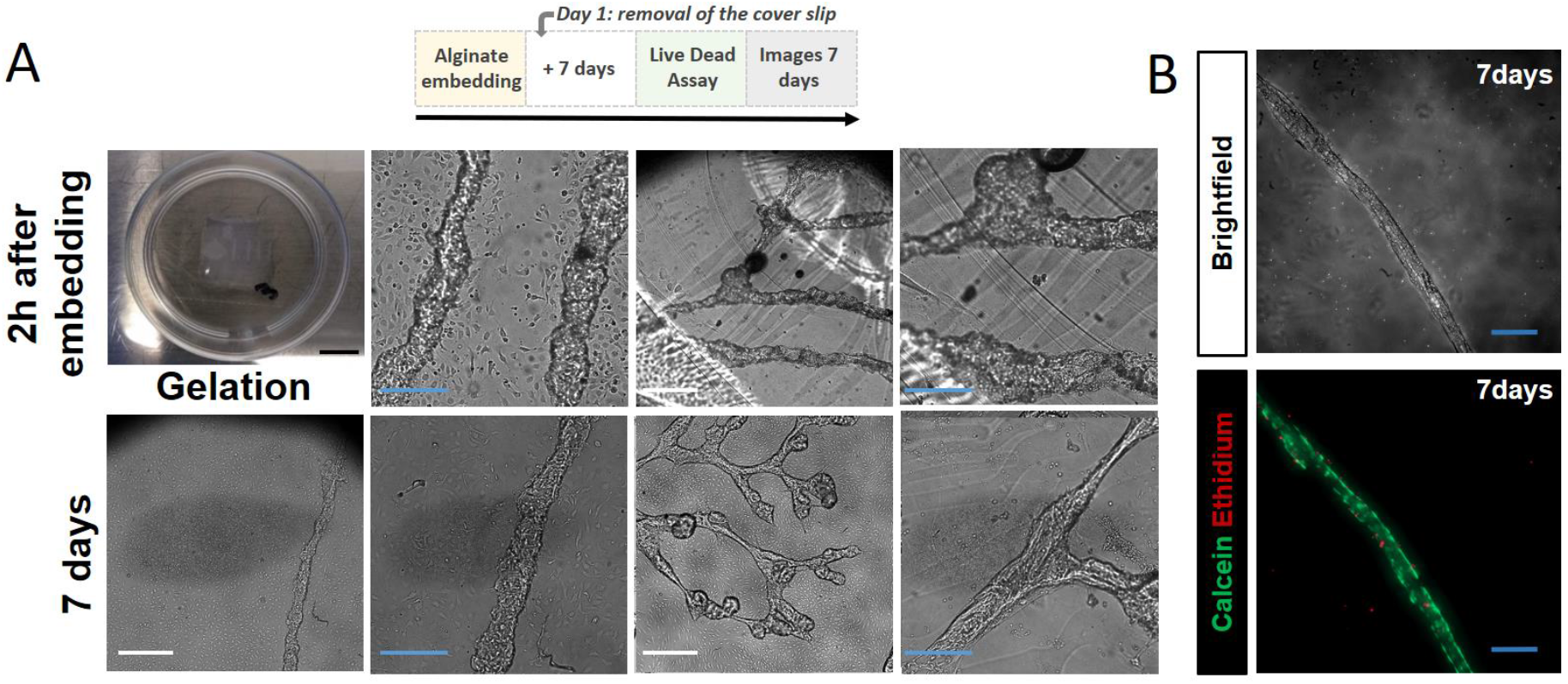
Images representing EPC structures (A) and cell viability (B), 7 days after embedding in an alginate patch. In the Live/Dead Assay (B), live cells are labelled by calcein (green) and dead cells by ethidium (red). Blue scale bars are 200 μm and white scale bars are 500 μm.

## CONCLUSION

Endothelial cells are known to spontaneously and rapidly form cord-like structures when plated in a basal membrane extract in medium containing angiogenic factors. This propensity is used as a functionality readout in what is known as an endothelial tube formation assay, a widely used method for studying in vitro angiogenesis. Tube formation is sustained for 18-24 h, and lumens may randomly form along the cords, after which time the tube networks disintegrate. The addition of pericytes^46^ or MSCs^47^ has been shown to stabilize the formation of endothelial luminal tubes in 3D co-culture conditions. We show here that not only can we create long-lasting networks of endothelial tubes from endothelial cells alone, but we can also build linear tubes spanning millimeters, as well as branched networks mimicking a microvascular bed according to a predetermined design. Hence, this work stands as a striking illustration of how engineering techniques, in particular gel micropatterning as well as bioprinting, can sustain the full potential of cellular self-organization into reproducibly building functional tissues and mini-organs with higher fidelity and robustness, as discussed previously in our review^10^, and demonstrated experimentally in recent advances by our team and other labs^26,37,38,48,49^. Compared to our previous study with biliary tubes^26^, endothelial cells appeared to require a stiffer environment to self-organize into stable tubes and we therefore had to resort to a more complex hydrogel mix associating both gelatin and collagen I, with hyaluronic acid and fibrin to reach a formulation that consistently showed efficiency in forming stable networks with EPCs from various donors. We also showed that this formulation showed excellent efficiency in promoting tube formation with a wide panel of other endothelial cell types (from both commercially-available HUVECs, HDMECs, as well as from HDLECs), thus proposing a global approach to create functional, robust, and reproducible in vitro vascular models which are much needed for a wide range of pharmacological and clinical applications.

Our approach shows in particular great potential in tissue engineering applications. Indeed, as no bulk hydrogel is present around the structures, the endothelial vessels obtained in this setup offer full access to the outside of the tube, allowing in follow-up studies the introduction of other cell types, such as smooth muscle cells and pericytes, at any stage of the vascular bed maturation, to obtain more complex and physiological blood vessel models. What’s more, by demonstrating that we can recover the structures in a soft patch and manipulate it as a sheet, we can envisage follow-up studies where we would integrate this sheet with other prefabricated tissues and organoids by essentially stacking the thin sheets one on top of the other. This however is no trivial feat as one is immediately confronted to issues of optimizing cell medium composition to accommodate the needs of all cell types that are present.

Perfusion of this vascular model *in vitro* remains however a challenge. Even though we show first successful attempts at microinjecting fluorescent dyes into the tubular network, developing a robust method to connect a microfluidic system to the inlet and outlet ports of such a network to bring nutrients and oxygen to the bioengineered tissue, and recapitulate physiological shear stress, still needs to be addressed. As the 4-component hydrogel formulation seems to partially degrade over time, it would also be interesting to elucidate if the cells are replacing this biodegradable scaffold with newly synthesized ECM, either on the apical side or basal side, and if the addition of pericytes may further contribute to the formation, maintenance, and remodeling of a basement membrane^46,50^.

## Supporting information

Supplemental movie Tree

## FUNDING AND ACKNOWLEDGMENTS

This work received the financial support of the iLite RHU program (grant ANR ANR-16-RHUS-0005) and from MSDAvenir via the Bio3DHE project.

We thank Didier Letourneur and Isabelle Bataille (INSERM U1148, Univ Paris 13) for access to their rheology equipment, the Technological Core Facility of the Institut de Recherche Saint-Louis, Université Paris-Cité for continued support in multimodal microscopy, Lousineh Arakelian (Team 7 INSERM U976) for assistance and expertise in EPC isolation, and the CytomorphoLab (Team 8 INSERM U976) for assistance on microinjection and access to their microfabrication room.

## AUTHOR CONTRIBUTIONS

E.M-A. D.A. M.L., F.C. and A.F. conceived and designed the experiments; E.M-A. D.A., and M.L. performed the experimental work and contributed to the manipulation of materials, reagents, and analysis tools; E.M-A. D.A. M.L., F.C., L.F., J.L, and A.F. analyzed the data and wrote the manuscript. All authors read and approved the final manuscript.

## DATA AVAILABILITY

The raw/processed data required to reproduce these findings cannot be shared on a permanent website at this time due to technical or time limitations. The raw/processed data required to reproduce these findings can be shared by the authors upon request.

## SUPPLEMENTARY DATA

Movies: 50u.avi & Tree.avi: Timelapse acquisitions of EPCs embedded in a line-shaped hydrogel pattern of 50μm width (50u.avi) or in a tree-shaped hydrogel pattern (Tree.Avi) over 60 hours in culture. Phase contrast images were captured every 30 min starting 24h after photopolymerization (day 0), therefore spanning day 1 to day 5.

---

**Figure S1:**
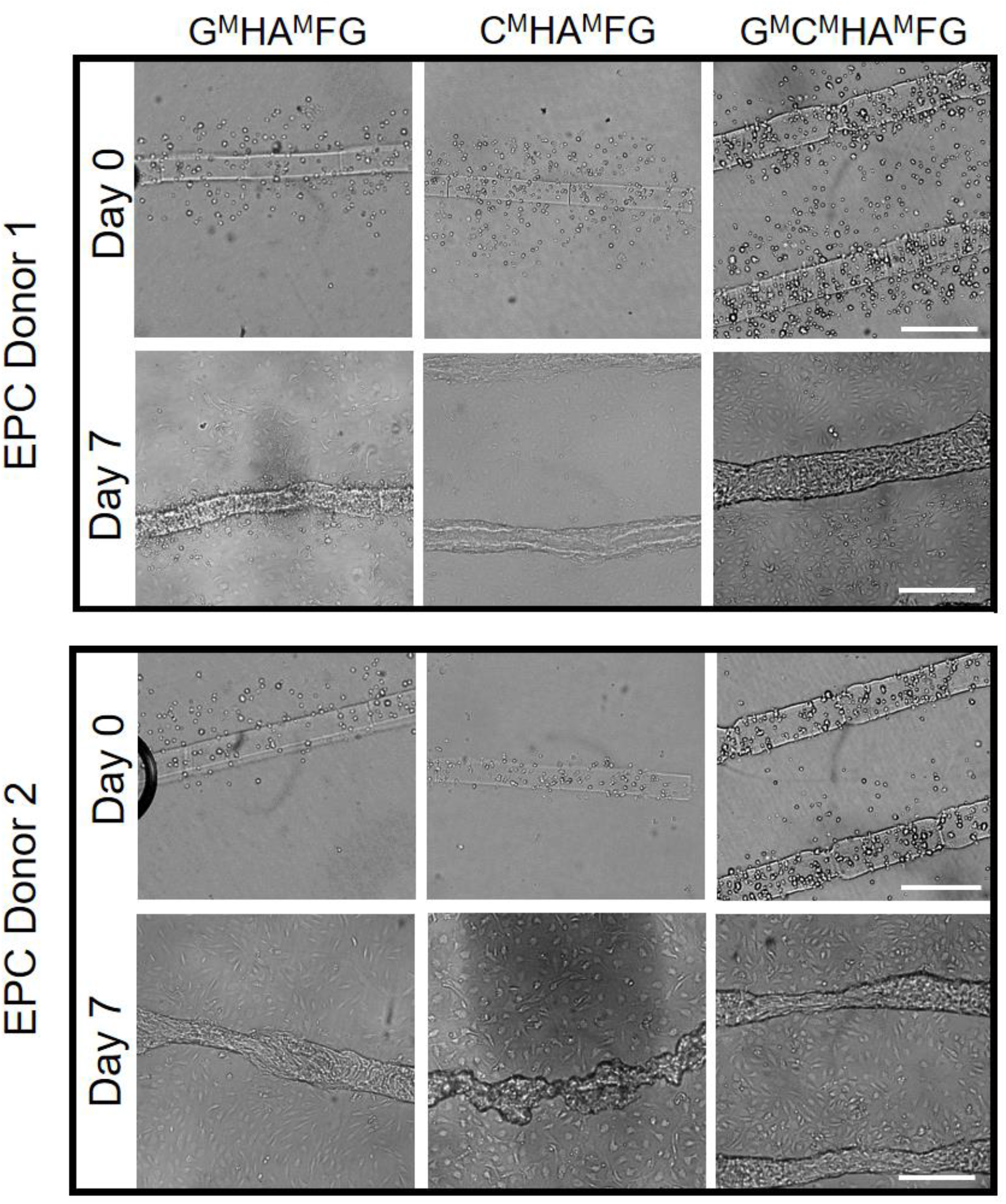
Phase contrast images showing structures obtained from EPCs isolated from 2 different donors that were encapsulated by photo-polymerization through a 10X objective at 20 mJ/mm^2^. Hydrogel formulations are indicated at the top. EPCs from donor 2 failed consistently to form stable tubuloid structures as seen in the lower middle image. Scale bars are 200 μm.

**Figure S2:**
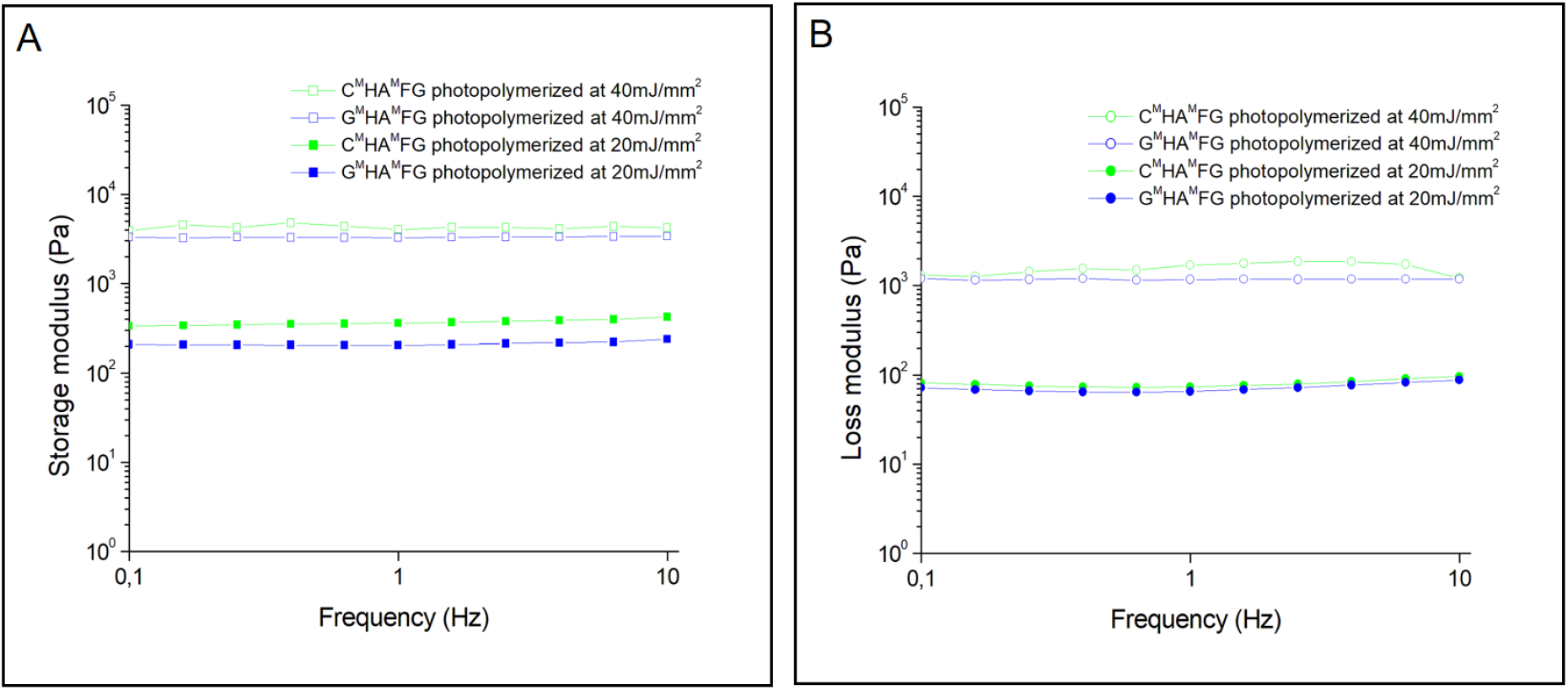
Rheological characterizations of 3-component photopolymerizes hydrogel formulations using frequency-dependent oscillatory rheological analysis as a function of UV dose. Resulting (A) storage moduli. (B) loss moduli.

**Figure S3.**
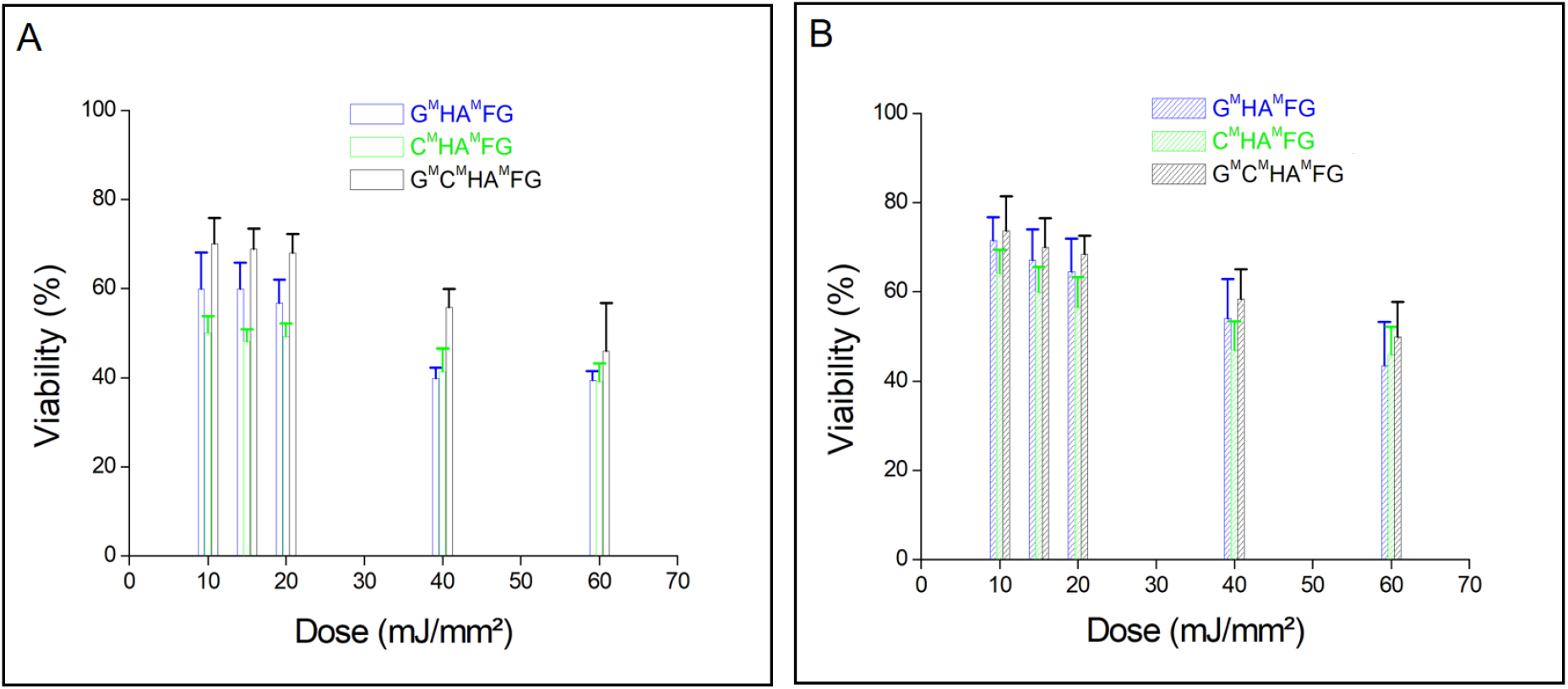
Cell viability of EPCs encapsulated within 3 different hydrogel formulations after photopolymerization. The bar chart showing the live cell percentage 4h (A) and 24h (B) after the photopolymerization process, measured using a standard live/dead assay. Error bars represent SEM and n = 4 for all data points. G^MA^/HA^MA^/FG hydrogels were mixed at 2%:0.3%:0.4% (w/v), C^MA^/HA^MA^/FG hydrogels were mixed at 0.2%:0.3%:0.4% (w/v), and G^MA^/C^MA^/HA^MA^/FG hydrogels were mixed at 1%:0.1%:0.3%:0.4% (w/v).

**Figure S4:**
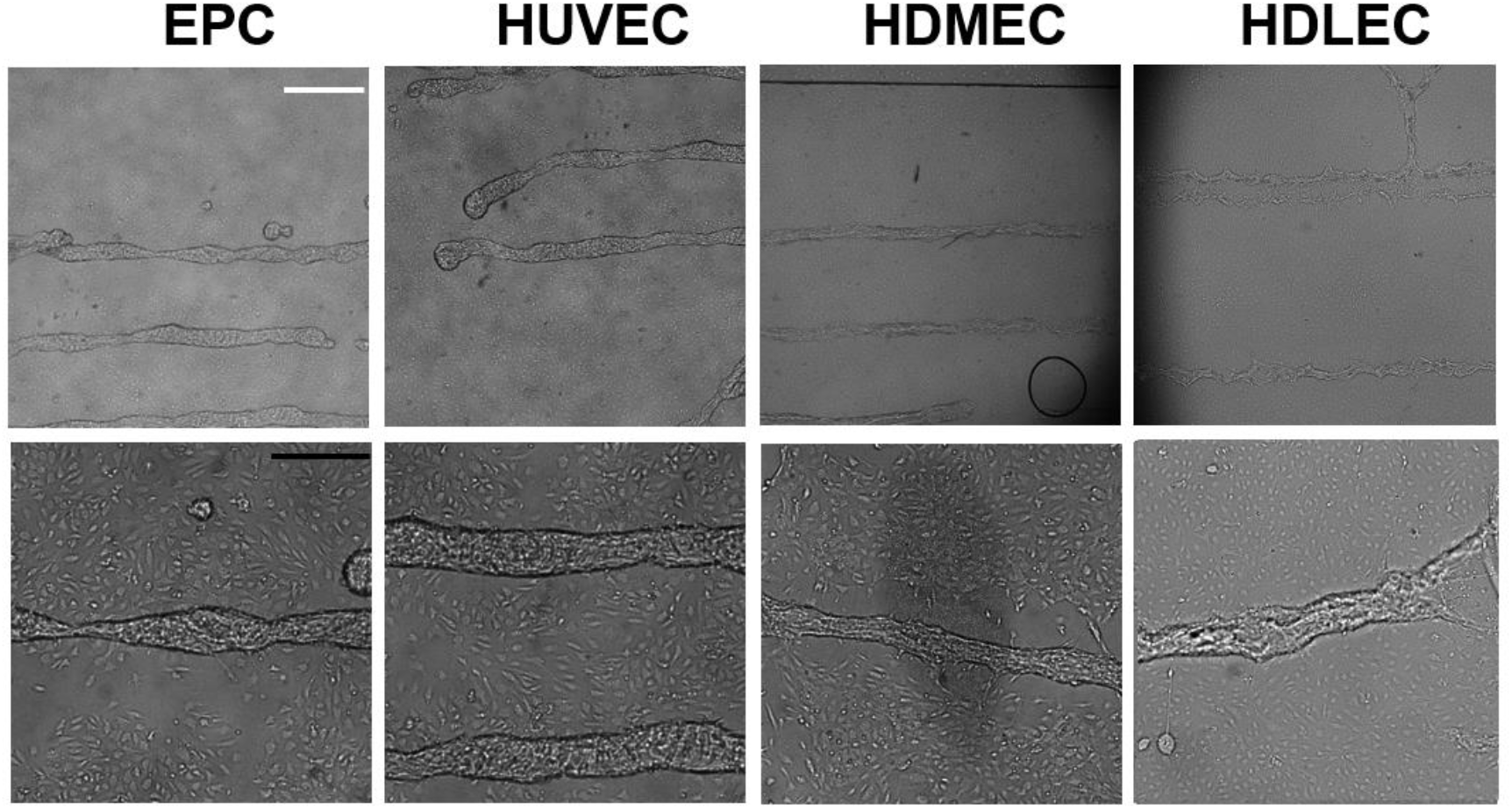
Culture of various sources of ECs embedded within a G^MA^/C^MA^/HA^MA^/FG 1%:0.1%:0.3%:0.4% (w/v) hydrogel formulation with 0.5% (wt/vol) photoinitiator, photo-polymerized through a 10X objective at 20 mJ/mm^2^. White scale bar is 500 μm and black scale bar is 200 μm.

## REFERENCES

1. Folkman, J. & Hochberg, M. Self-regulation of Growth in Three Dimensions. Journal of Experimental Medicine 138, 745–753 (1973).

2. Wang, Z., Mithieux, S. M. & Weiss, A. S. Fabrication Techniques for Vascular and Vascularized Tissue Engineering. Advanced Healthcare Materials 8, 1900742 (2019).

3. Ahlmann, A. H. et al. Decellularised Human Umbilical Artery as a Vascular Graft Elicits Minimal Pro-Inflammatory Host Response Ex Vivo and In Vivo. International Journal of Molecular Sciences 22, 7981 (2021).

4. Sulaiman, N. S. et al. Effective decellularisation of human saphenous veins for biocompatible arterial tissue engineering applications: Bench optimisation and feasibility in vivo testing. J Tissue Eng 12, 2041731420987529 (2021).

5. Wimmer, R. A., Leopoldi, A., Aichinger, M., Kerjaschki, D. & Penninger, J. M. Generation of blood vessel organoids from human pluripotent stem cells. Nat Protoc 14, 3082–3100 (2019).

6. Homan, K. A. et al. Flow-enhanced vascularization and maturation of kidney organoids in vitro. Nature Methods 1 (2019) doi:10.1038/s41592-019-0325-y.

7. Ebner-Peking, P. et al. Self-assembly of differentiated progenitor cells facilitates spheroid human skin organoid formation and planar skin regeneration. Theranostics 11, 8430–8447 (2021).

8. Ahn, Y. et al. Human Blood Vessel Organoids Penetrate Human Cerebral Organoids and Form a Vessel-Like System. Cells 10, 2036 (2021).

9. Wimmer, R. A. et al. Human blood vessel organoids as a model of diabetic vasculopathy. Nature 565, 505–510 (2019).

10. Laurent, J. et al. Convergence of microengineering and cellular self-organization towards functional tissue manufacturing. Nature Biomedical Engineering 1, 939–956 (2017).

11. Rim, N. G. et al. Micropatterned cell sheets as structural building blocks for biomimetic vascular patches. Biomaterials 181, 126–139 (2018).

12. Negishi, J. et al. Evaluation of small-diameter vascular grafts reconstructed from decellularized aorta sheets. Journal of Biomedical Materials Research Part A 105, 1293–1298 (2017).

13. He, W., Sharma, D., Jia, W. & Zhao, F. Fabrication of a Completely Biological and Anisotropic Human Mesenchymal Stem Cell-Based Vascular GraftVascular grafts. in Vascular Tissue Engineering: Methods and Protocols (eds. Zhao, F. & Leong, K. W.) 101–114 (Springer US, 2022). doi:10.1007/978-1-0716-1708-3_9.

14. Millik, S. C. et al. 3D printed coaxial nozzles for the extrusion of hydrogel tubes toward modeling vascular endothelium. Biofabrication 11, 045009 (2019).

15. Patel, H. N., Vohra, Y. K., Singh, R. K. & Thomas, V. HuBiogel incorporated fibro-porous hybrid nanomatrix graft for vascular tissue interfaces. Materials Today Chemistry 17, 100323 (2020).

16. Nguyen, T.-U., Shojaee, M., Bashur, C. A. & Kishore, V. Electrochemical fabrication of a biomimetic elastin-containing bi-layered scaffold for vascular tissue engineering. Biofabrication 11, 015007 (2018).

17. Zheng, F., Derby, B. & Wong, J. Fabrication of microvascular constructs using high resolution electrohydrodynamic inkjet printing. Biofabrication 13, 035006 (2021).

18. Polacheck, W. J., Kutys, M. L., Tefft, J. B. & Chen, C. S. Microfabricated blood vessels for modeling the vascular transport barrier. Nat Protoc 14, 1425–1454 (2019).

19. Yuan, B. et al. A Strategy for Depositing Different Types of Cells in Three Dimensions to Mimic Tubular Structures in Tissues. Advanced Materials 24, 890–896 (2012).

20. Bosch-Rué, E., Delgado, L. M., Gil, F. J. & Perez, R. A. Direct extrusion of individually encapsulated endothelial and smooth muscle cells mimicking blood vessel structures and vascular native cell alignment. Biofabrication 13, 015003 (2020).

21. Norouzi, S. K. & Shamloo, A. Bilayered heparinized vascular graft fabricated by combining electrospinning and freeze drying methods. Materials Science and Engineering: C 94, 1067–1076 (2019).

22. Carmeliet, P. & Jain, R. K. Angiogenesis in cancer and other diseases. Nature 407, 249–257 (2000).

23. Burr, S. et al. Oxygen gradients can determine epigenetic asymmetry and cellular differentiation via differential regulation of Tet activity in embryonic stem cells. Nucleic Acids Research 46, 1210–1226 (2018).

24. Zhang, A. P. et al. Rapid Fabrication of Complex 3D Extracellular Microenvironments by Dynamic Optical Projection Stereolithography. Adv. Mater. 24, 4266–4270 (2012).

25. Nicolas, J. et al. 3D Extracellular Matrix Mimics: Fundamental Concepts and Role of Materials Chemistry to Influence Stem Cell Fate. Biomacromolecules 21, 1968–1994 (2020).

26. Mazari-Arrighi, E. et al. Construction of functional biliary epithelial branched networks with predefined geometry using digital light stereolithography. Biomaterials 279, 121207 (2021).

27. Wu, L. et al. Protocol update for late endothelial progenitor cells identified by double-positive staining. Journal of Cellular and Molecular Medicine 26, 306–311 (2022).

28. Lafleur, M. A., Handsley, M. M., Knäuper, V., Murphy, G. & Edwards, D. R. Endothelial tubulogenesis within fibrin gels specifically requires the activity of membrane-type-matrix metalloproteinases (MT-MMPs). J Cell Sci 115, 3427–3438 (2002).

29. Vernon, R. B. & Sage, E. H. A Novel, Quantitative Model for Study of Endothelial Cell Migration and Sprout Formation within Three-Dimensional Collagen Matrices. Microvascular Research 57, 118–133 (1999).

30. Hiraoka, N., Allen, E., Apel, I. J., Gyetko, M. R. & Weiss, S. J. Matrix metalloproteinases regulate neovascularization by acting as pericellular fibrinolysins. Cell 95, 365–377 (1998).

31. Ilan, N., Mahooti, S. & Madri, J. A. Distinct signal transduction pathways are utilized during the tube formation and survival phases of in vitro angiogenesis. Journal of Cell Science 111, 3621–3631 (1998).

32. Camci-Unal, G., Cuttica, D., Annabi, N., Demarchi, D. & Khademhosseini, A. Synthesis and Characterization of Hybrid Hyaluronic Acid-Gelatin Hydrogels. Biomacromolecules 14, 1085–1092 (2013).

33. Fairbanks, B. D., Schwartz, M. P., Bowman, C. N. & Anseth, K. S. Photoinitiated polymerization of PEG-diacrylate with lithium phenyl-2,4,6-trimethylbenzoylphosphinate: polymerization rate and cytocompatibility. Biomaterials 30, 6702–6707 (2009).

34. Gaudet, I. D. & Shreiber, D. I. Characterization of Methacrylated Type-I Collagen as a Dynamic, Photoactive Hydrogel. Biointerphases 7, 25 (2012).

35. Antoine, E. E., Vlachos, P. P. & Rylander, M. N. Review of Collagen I Hydrogels for Bioengineered Tissue Microenvironments: Characterization of Mechanics, Structure, and Transport. Tissue Eng Part B Rev 20, 683–696 (2014).

36. Bupphathong, S. et al. Gelatin Methacrylate Hydrogel for Tissue Engineering Applications-A Review on Material Modifications. Pharmaceuticals (Basel) 15, 171 (2022).

37. Brassard, J. A., Nikolaev, M., Hübscher, T., Hofer, M. & Lutolf, M. P. Recapitulating macro-scale tissue self-organization through organoid bioprinting. Nat. Mater. 20, 22–29 (2021).

38. Ma, X. et al. Deterministically patterned biomimetic human iPSC-derived hepatic model via rapid 3D bioprinting. PNAS 113, 2206–2211 (2016).

39. Stratesteffen, H. et al. GelMA-collagen blends enable drop-on-demand 3D printablility and promote angiogenesis. Biofabrication 9, 045002 (2017).

40. Davidov, T., Efraim, Y., Dahan, N., Baruch, L. & Machluf, M. Porcine arterial ECM hydrogel: Designing an in vitro angiogenesis model for long-term high-throughput research. The FASEB Journal 34, 7745–7758 (2020).

41. Kobayashi, M. et al. Elastic Modulus of ECM Hydrogels Derived from Decellularized Tissue Affects Capillary Network Formation in Endothelial Cells. International Journal of Molecular Sciences 21, 6304 (2020).

42. Kristofik, N. J. et al. Improving in vivo outcomes of decellularized vascular grafts via incorporation of a novel extracellular matrix. Biomaterials 141, 63–73 (2017).

43. Kong, H. J., Smith, M. K. & Mooney, D. J. Designing alginate hydrogels to maintain viability of immobilized cells. Biomaterials 24, 4023–4029 (2003).

44. Andersen, T., Auk-Emblem, P. & Dornish, M. 3D Cell Culture in Alginate Hydrogels. Microarrays (Basel) 4, 133–161 (2015).

45. Roche, C. D. et al. Printability, Durability, Contractility and Vascular Network Formation in 3D Bioprinted Cardiac Endothelial Cells Using Alginate-Gelatin Hydrogels. Frontiers in Bioengineering and Biotechnology 9, (2021).

46. Stratman, A. N. & Davis, G. E. Endothelial Cell-Pericyte Interactions Stimulate Basement Membrane Matrix Assembly: Influence on Vascular Tube Remodeling, Maturation, and Stabilization. Microscopy and Microanalysis 18, 68–80 (2012).

47. Duttenhoefer, F. et al. 3D scaffolds co-seeded with human endothelial progenitor and mesenchymal stem cells: evidence of prevascularisation within 7 days. Eur Cell Mater 26, 49–64; discussion 64-65 (2013).

48. Gontran, E. et al. Self-Organogenesis from 2D Micropatterns to 3D Biomimetic Biliary Trees. Bioengineering 8, 112 (2021).

49. Zhu, W. et al. Direct 3D bioprinting of prevascularized tissue constructs with complex microarchitecture. Biomaterials 124, 106–115 (2017).

50. Geevarghese, A. & Herman, I. M. Pericyte-Endothelial Cross-Talk: Implications and Opportunities for Advanced Cellular Therapies. Transl Res 163, 296–306 (2014).

